# Chemical composition and the potential for proteomic transformation in cancer, hypoxia, and hyperosmotic stress

**DOI:** 10.1101/097667

**Authors:** Jeffrey M. Dick

## Abstract

Models of chemical composition and thermodynamic potential emphasize the microenvironmental context for proteomic transformations. Here, data from 71 comparative proteomics studies were analyzed using elemental ratios and chemical components as measures of oxidation and hydration state. Experimental lowering of oxygen availability (hypoxia) and water activity (hyperosmotic stress) are reflected in decreased oxidation and hydration states of proteomes, but up-expressed proteins in colorectal and pancreatic cancer most often have higher average oxidation or hydration state. Calculations of chemical affinity were used to quantify the thermodynamic potentials for transformation of proteomes as a function of fugacity of O_2_ and activity of H_2_O, which serve as scales of oxidation and hydration potential. Potential diagrams for the overall reactions between down- and up-expressed proteins in various datasets have predicted equipotential lines that cluster near unit activity of H_2_O, suggesting that protein expression is sensitive to local variations in water activity. A redox balance calculation indicates that an increase in the lipid to protein ratio in cancer cells by 20% over hypoxic cells would generate an electron sink of sufficient size for oxidation of the cancer proteomes. These findings demonstrate consistent, condition-dependent chemical differences between proteomes, identify primary microenvironmental constraints on proteomic transformations, and help to quantify the redox balance among cellular macromolecules. The datasets and computer code used here are supplied in a new R package, **canprot**.

## INTRODUCTION

The relationship between cells and tissue microenvironments is a topic of vital importance for cancer biology. Because of rapid cellular proliferation and irregular vascularization, tumors often develop regions of hypoxia (Höckel and Vaupel, 2001). Tumor microenvironments also exhibit abnormal ranges for other physical-chemical variables, including hydration state (McIntyre, 2006; Abramczyk et al., 2014).

Some aspects of the complex metazoan response to hypoxia are mediated by hypoxia-inducible factor 1 (HIF-1). HIF-1 is a transcription factor that is tagged for degradation in normoxic conditions. Under hypoxia, the degradation of HIF-1 is suppressed; HIF-1 can then enter the nucleus and activate the transcription of downstream targets (Semenza, 2003). Indeed, transcriptional targets of HIF-1 are found to be differentially expressed in proteomic datasets for laboratory hypoxia (Cifani et al., 2011; McMahon et al., 2012). However, proteomic studies of hypoxia provide many examples of proteins that are not directly regulated by HIF-1 (McMahon et al., 2012; Fuhrmann et al., 2013), and cancer proteomic datasets also include many proteins that are not known to be regulated by HIF-1.

The complexity of underlying regulatory mechanisms (McMahon et al., 2012) combined with extensive dissimilarity between gene transcription and protein translation (van den Beucken et al., 2011; Cifani et al., 2011; Ho et al., 2016) present many difficulties for a bottom-up understanding of global proteomic trends. As a counterpart to molecular explanations, a systems perspective can incorporate higher-level constraints (Drack and Wolkenhauer, 2011). A commonly used metaphor in systems biology is attractor landscapes. The basins of attraction are defined by dynamical systems behavior, but are also analogous to minimum-energy states in thermodynamics (Emmeche et al., 2000; Enver et al., 2009). Nevertheless, few studies have investigated the thermodynamic potentials that are inherent in the compositional biology of proteomic transformations.

To better understand the microenvironmental context for compositional changes, this paper uses proteomic data as input into a high-level thermodynamic model. First, a compositional analysis of differentially (down- and up-) expressed proteins identifies consistent trends in the oxidation and hydration states of proteomes of colorectal cancer (CRC), pancreatic cancer, and cells exposed to hypoxia or hyperosmotic stress. These results lay the groundwork for using a thermodynamic model to quantify environmental constraints on the potential for proteomic transformation. Finally, the Discussion explores some implications of the hypothesis that elevated synthesis of lipids provides an electron sink for the oxidation of proteomes. In this view, some cancer systems may develop an abnormally large redox disproportionation among pools of cellular biomacromolecules.

## METHODS

### Data sources

Tables 1-4 present the sources of data. Protein IDs and expression (up/down or abundance ratios) were found in the literature, often being reported in the supporting information (SI) or supplementary (suppl.) tables. In some cases, source tables were further processed, using fold-change and significance cutoffs that, where possible, are based on statements made in the primary publication. The data are stored as *.csv files in the R package **canprot**, which was developed during this study (see http://github.com/jedick/canprot) and is provided as Dataset S1.

**Table 1.**
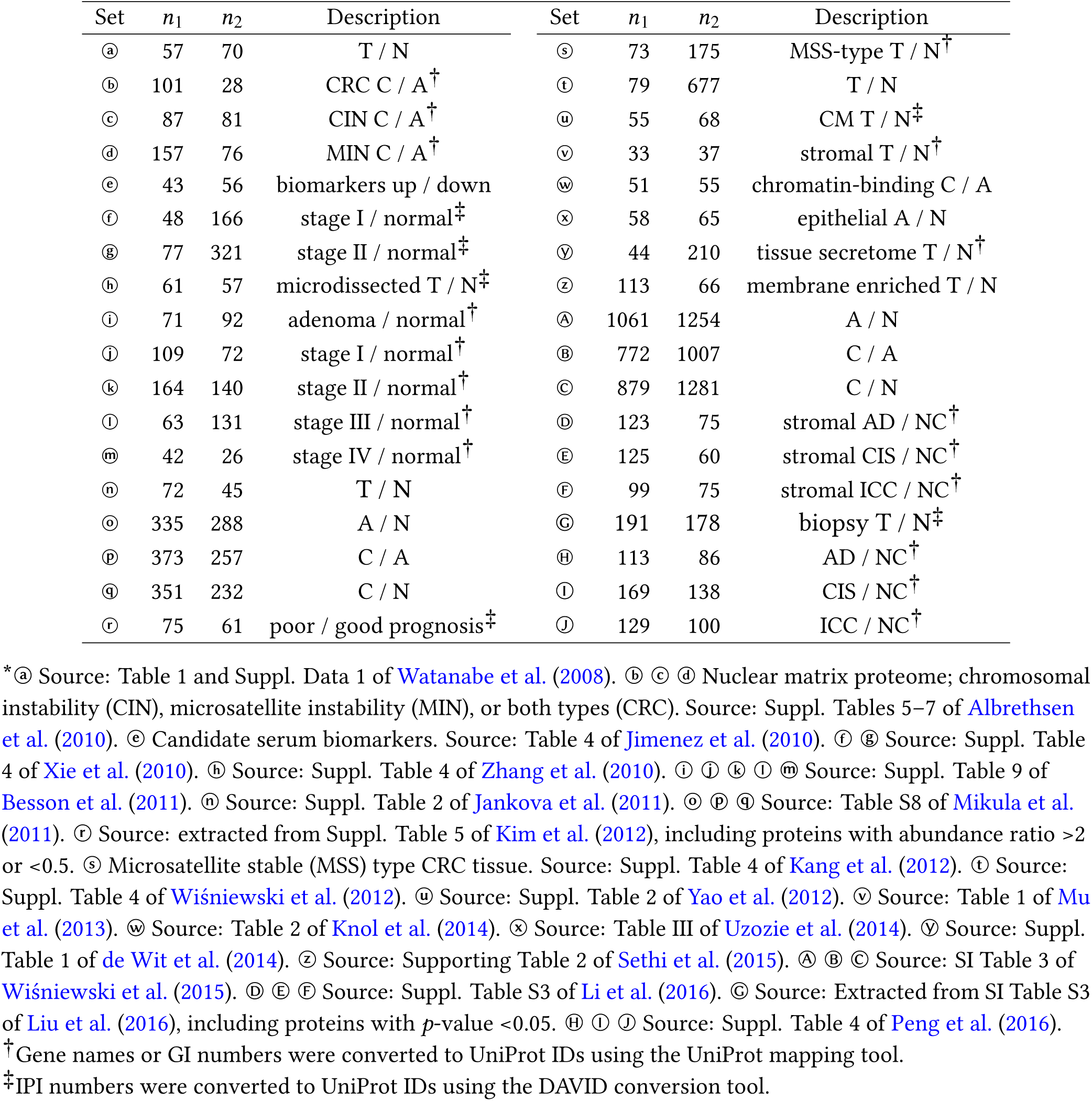
Selected proteomic datasets for colorectal cancer.* Abbreviations: T (tumor), N (normal), C (carcinoma or adenocarcinoma), A (adenoma), CM (conditioned media), AD (adenomatous colon polyps), CIS (carcinoma *in situ),* ICC (invasive colonic carcinoma), NC (non-neoplastic colonic mucosa).

**Table 2.**
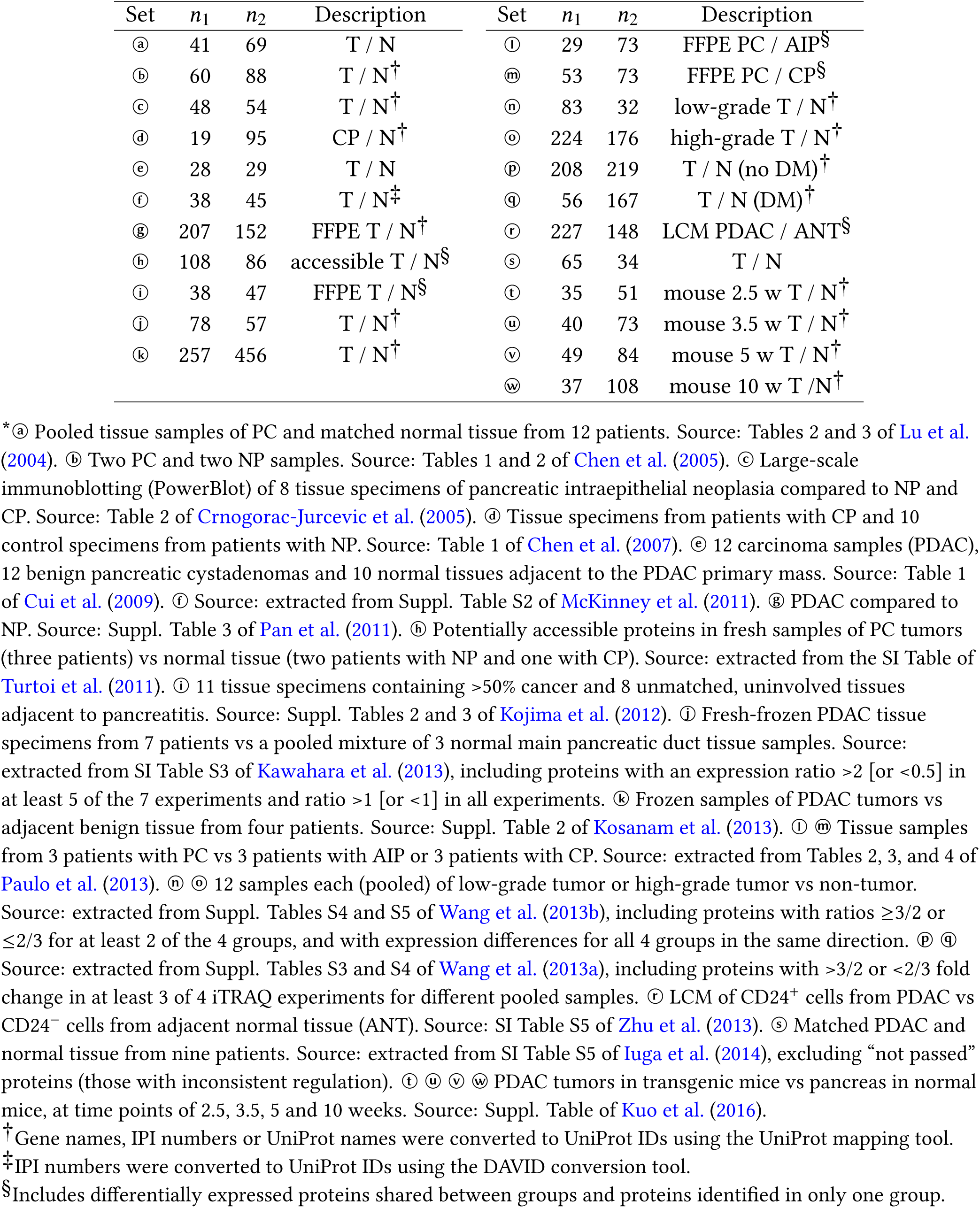
Selected proteomic datasets for pancreatic cancer.* Abbreviations: T (tumor), N (normal), CP (chronic pancreatitis), AIP (autoimmune pancreatitis), PC (pancreatic cancer), DM (diabetes mellitus), PDAC (pancreatic ductal adenocarcinoma), ANT (adjacent normal tissue), FFPE (formalin-fixed paraffin-embedded), LCM (laser-capture microdissection), NP (normal pancreas).

**Table 3.**
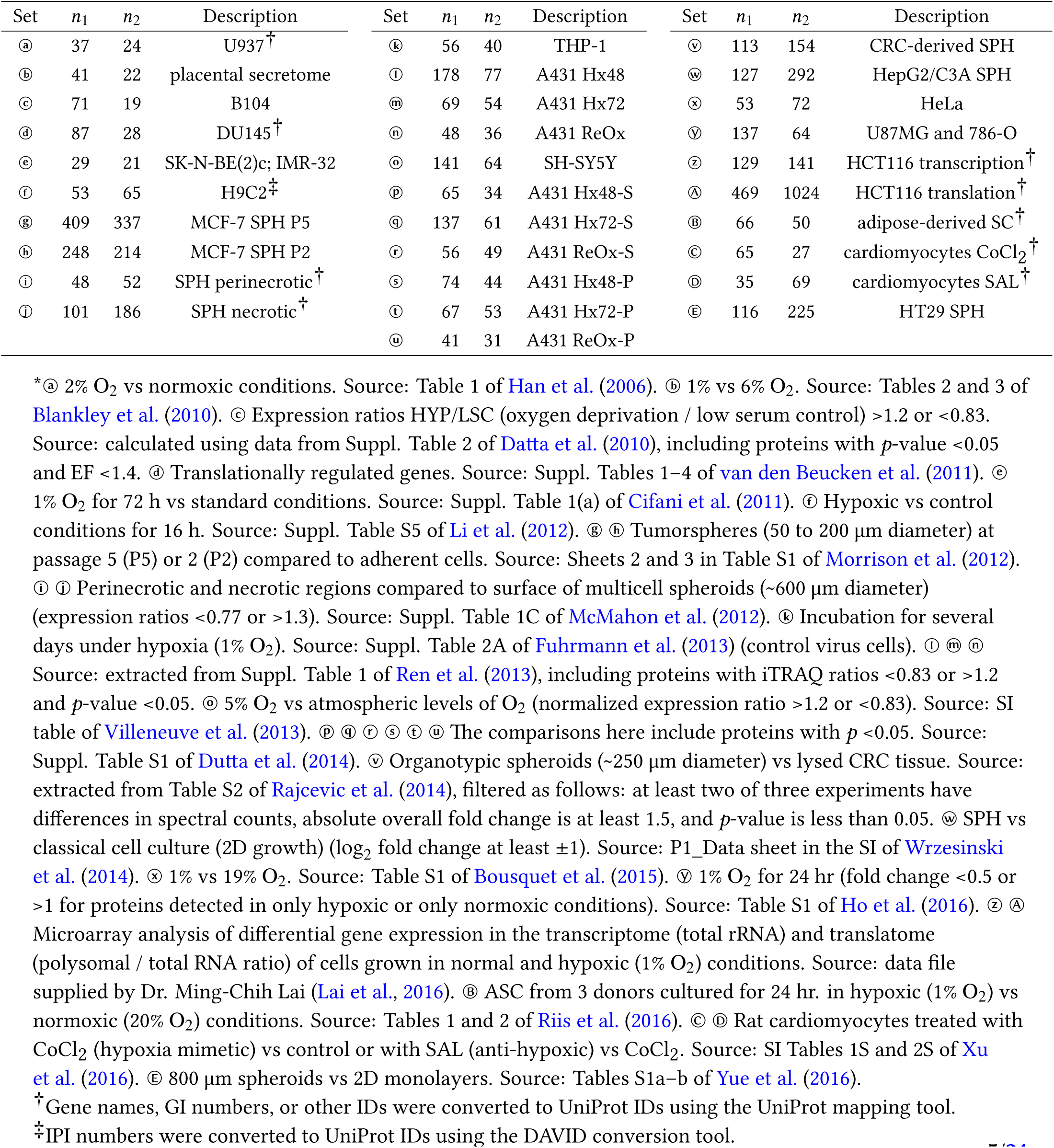
Selected proteomic datasets for hypoxia and reoxygenation experiments.* Abbreviations: U937 (acute promonocytic leukemic cells), B104 (rat neuroblastoma cells), DU145 (prostate carcinoma cells), SK-N-BE(2)c; IMR-32; SH-SY5Y (neuroblastoma cells), H9C2 (rat heart myoblast), MCF-7 (breast cancer cells), THP-1 (macrophages), A431 (epithelial carcinoma cells), Hx48 (hypoxia 48 h), Hx72 (hypoxia 72 h), ReOx (hypoxia 48 h followed by reoxygenation for 24 h), -S (supernatant fraction), -P (pellet fraction), SPH (spheroids), HepG2/C3A (hepatocellular carcinoma cells), U87MG (glioblastoma), 786-O (renal clear cell carcinoma cells), HCT116; HT29 (colon cancer cells), SC (stem cells), SAL (salidroside).

**Table 4.**
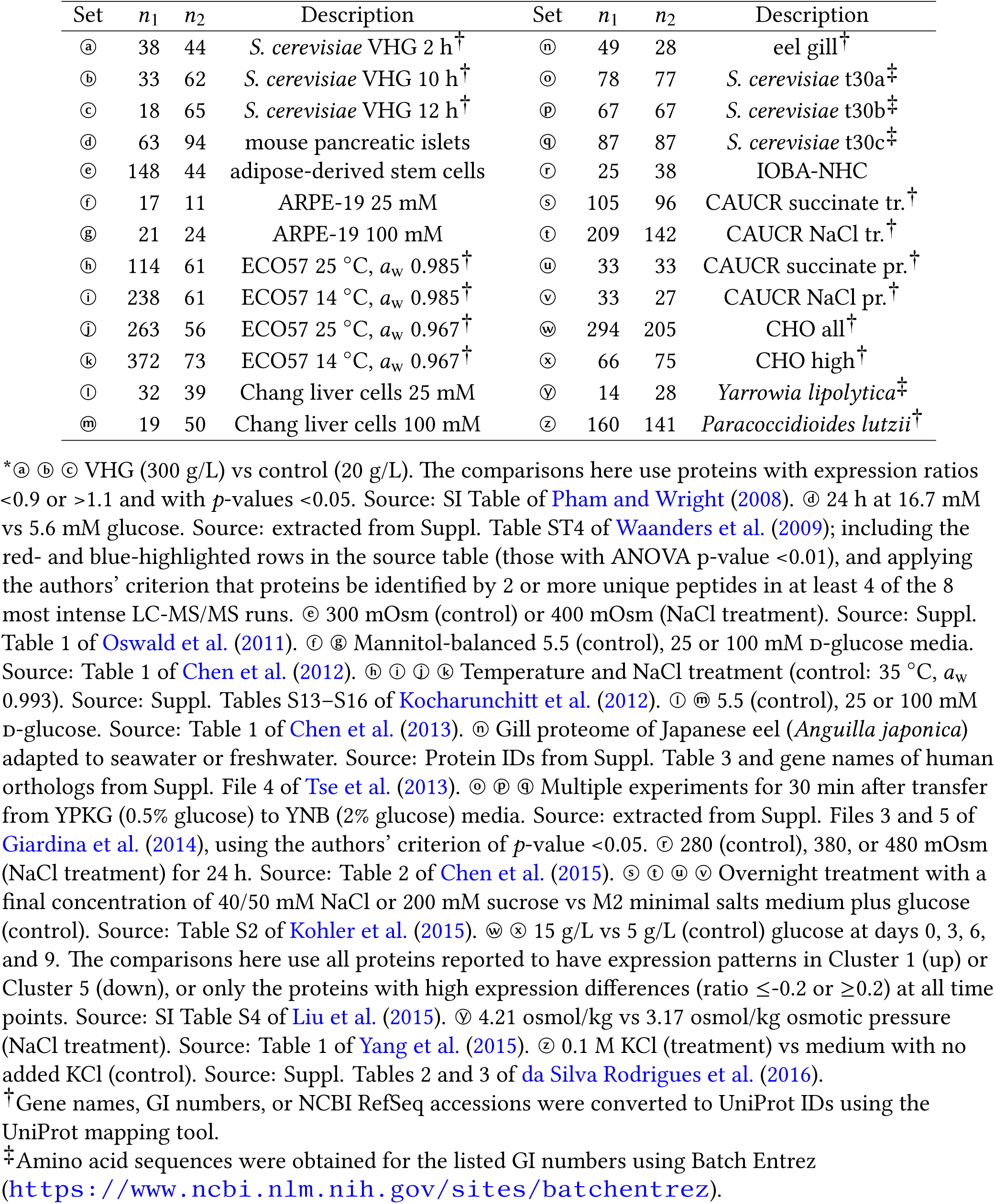
Selected proteomic datasets for hyperosmotic stress experiments.* Abbreviations: VHG (very high glucose), ARPE-19 (human retinal pigmented epithelium cells), ECO57 *(Escherichia coli* O157:H7 Sakai), IOBA-NHC (human conjunctival epithelial cells), CAUCR *(Caulobacter crescentus),* tr. (transcriptome), pr. (proteome), CHO (Chinese hamster ovary cells).

Sequence IDs were converted to UniProt IDs using the UniProt mapping tool (http://www.uniprot.org/mapping/) or the gene ID conversion tool of DAVID 6.7 (https://david.ncifcrf.gov/conversion.jsp). For proteins where the automatic conversions produced no matches, manual searches in UniProt were performed using the gene names or protein descriptions. If specified (i.e. as UniProt IDs with suffixes), particular isoforms of the proteins were used.

Obsolete or secondary IDs reported for some proteins were updated to reflect current, primary IDs (uniprot_updates.csv in Dataset S1). Any duplicated IDs listed as having opposite expression ratios were excluded from the comparisons here.

Amino acid sequences of human proteins were taken from the UniProt human reference proteome. Sequences of proteins in other organisms and of human proteins not contained in the reference proteome were downloaded from UniProt or other sources. Amino acid compositions were computed using functions in the CHNOSZ package (Dick, 2008) or the ProtParam tool on the UniProt website (see Dick, 2016). The amino acid compositions are stored in *.csv files in Dataset S1.

R (R Core Team, 2016) and R packages **canprot** (this study) and **CHNOSZ** (Dick, 2008) were used to process the data and generate the figures with code specifically written for this study, which is provided in Dataset S2.

### Compositional metrics for oxidation and hydration state

Two compositional metrics that afford a quantitative description of proteomic data, the average oxidation state of carbon (*Z*_c_) and the water demand per residue (*n*̅H_2_O), are briefly described here.

The oxidation state of atoms in molecules quantifies the degree of electron redistribution due to bonding; a higher oxidation state signifies a lower degree of reduction. Although calculations of oxidation state from molecular formulas necessarily make simplifying assumptions regarding the internal electronic structure of molecules, such calculations may be used to quantify the flow of electrons in chemical reactions, and the oxidation state concept is useful for studying the transformations of complex mixtures of organic molecules. For example, calculations of the average oxidation state of carbon provide insight on the processes affecting decomposition of carbohydrate, protein and lipid fractions of natural organic matter (Baldock et al., 2004). Moreover, oxidation state can be regarded as an ensemble property of organic systems (Kroll et al., 2015). See Dick, 2016 for additional references where organic and biochemical reactions have been characterized using the average oxidation state of carbon.

Despite the large size of proteins, their relatively simple primary structure means that the *Z*_c_ can be computed using the elemental abundances in any particular amino acid sequence (Dick, 2014):

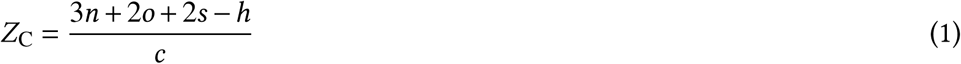

In this equation, *c*, *h*, *n*, *o*, and *s* are the elemental abundances in the chemical formula C_*c*_H_*h*_N_*n*_O_*O*_S_*s*_ for a specific protein.

In contrast to the elemental stoichiometry in Eq. (1), a calculation of the hydration state must take account of the gain or loss of H_2_O. In the biochemical literature, “protein hydration” or water of hydration refers to the effective (time-averaged) number of water molecules that interact with a protein (Timasheff, 2002). Protein hydration has important implications for crystallography and enzymatic function, but hydration numbers have been measured for few proteins and are difficult to compute, especially for the many proteins with unknown tertiary structure. Thus, the structural hydration of proteins identified in proteomic datasets generally remains unquantified.

A different concept of hydration state arises by considering the chemical components that make up proteins. A componential analysis is a method of projecting the composition of a molecule using specified chemical formula units as the components, or basis species. However, the implications go beyond mathematical considerations. The notion of components is central to chemical thermodynamics (Gibbs, 1875); the choice of components determines the thermodynamic variables (chemical potentials), and a careful choice leads to more convenient representations of the compositional and energetic constraints on reactions (e.g. Zhu and Anderson, 2002).

Dick (2016) proposed using C_5_H_10_N_2_O_3_, C_5_H_9_NO_4_, C_3_H_7_N_O2_S,O_2_, and H_2_O as a basis for assessing compositional differences in proteomes. The first three formulas correspond to glutamine, glutamic acid, and cysteine. Compared to common inorganic species, this basis reduces the projected codependence of oxidation and hydration state in proteins, effectively unfolding a compositional dimension that can enrich a thermodynamic model. Using this basis, the overall formation of a protein with formula C_*c*_H_*h*_N_*n*_O_*O*_S_*s*_ is represented by

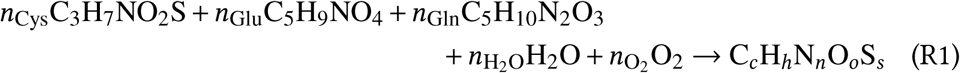

This stoichiometric reaction is not a mechanism for protein formation, rather a projection of any protein’s elemental composition into chemical components, i.e. the basis. To compare the compositions of different-sized proteins, the stoichiometric coefficients in Reaction R1 are divided by the respective sequence lengths (number of amino acids) of the proteins. The length-normalized coefficients, written with an overbar, include the per-residue water demand for formation of a protein (*n*̅H_2_O).This componential “hydration state” is used in this study, and should not be confused with the structural biochemical “protein hydration” mentioned above.

## RESULTS

### Colorectal cancer

The progression of colorectal cancer (CRC) begins with the formation of numerous non-cancerous lesions (adenomas), which may remain undetectable. Over time, a small fraction of adenomas develop into malignant tumors (carcinoma). Publicly available datasets reporting a minimum of ca. 30 up- and 30 down-expressed proteins for tissue samples of CRC, and one meta-analysis of serum biomarkers, were compiled recently (Dick, 2016). These same datasets are listed in Table 1, with one newer addition (dataset Ⓖ; Liu et al., 2016).

Many aspects of the experimental methods, statistical tests, and bioinformatics analyses used to identify significantly down-expressed and up-expressed proteins vary considerably among studies. The comparisons here are made without any control of this variability. Although particular comparisons may reflect study-specific conditions, visualization of the chemical compositions of proteins for many datasets can reveal general attributes of the cancer phenotype.

For each dataset, Table 1 lists the numbers of down-expressed (*n*_1_) and up-expressed (*n*_2_) proteins in cancer relative to normal tissue. Mean values of average oxidation state of carbon (*Z*_C_; Eq. 1) and water demand per residue (*n*̅H_2_O; Reaction R1) were calculated for the down- and up-expressed groups of proteins, together with the corresponding mean differences Δ*Z*_C_ and Δ*n*̅H_2_O), *p*-values, and effect sizes. These values are listed in Table S1.

Fig. 1A shows Δ*n*̅H_2_O vs Δ*Z*_c_ for the CRC datasets. The gray boxes cover the range from −0.01 to 0.01 for each of the variables. To draw attention to the largest and most significant changes, filled points indicate mean differences with a *p*-value (Wilcoxon test) less than 0.05, and solid lines indicate mean differences with a percent value of common language effect size (CLES) ≥60% or ≤40%. The common language statistic “is the probability that a score sampled at random from one distribution will be greater than a score sampled from some other distribution” (McGraw and Wong, 1992). Here, CLES is calculated as the percentage of pairings of individual proteins with a positive difference in *Z*_C_ or *n*̅H_2_O between group 1 and group 2, among all possible pairings between groups. Point symbols are squares if the *p*-values for both *Z*_C_ and *n*̅H_2_O are less than 0.05, or circles otherwise.

**Figure 1.**
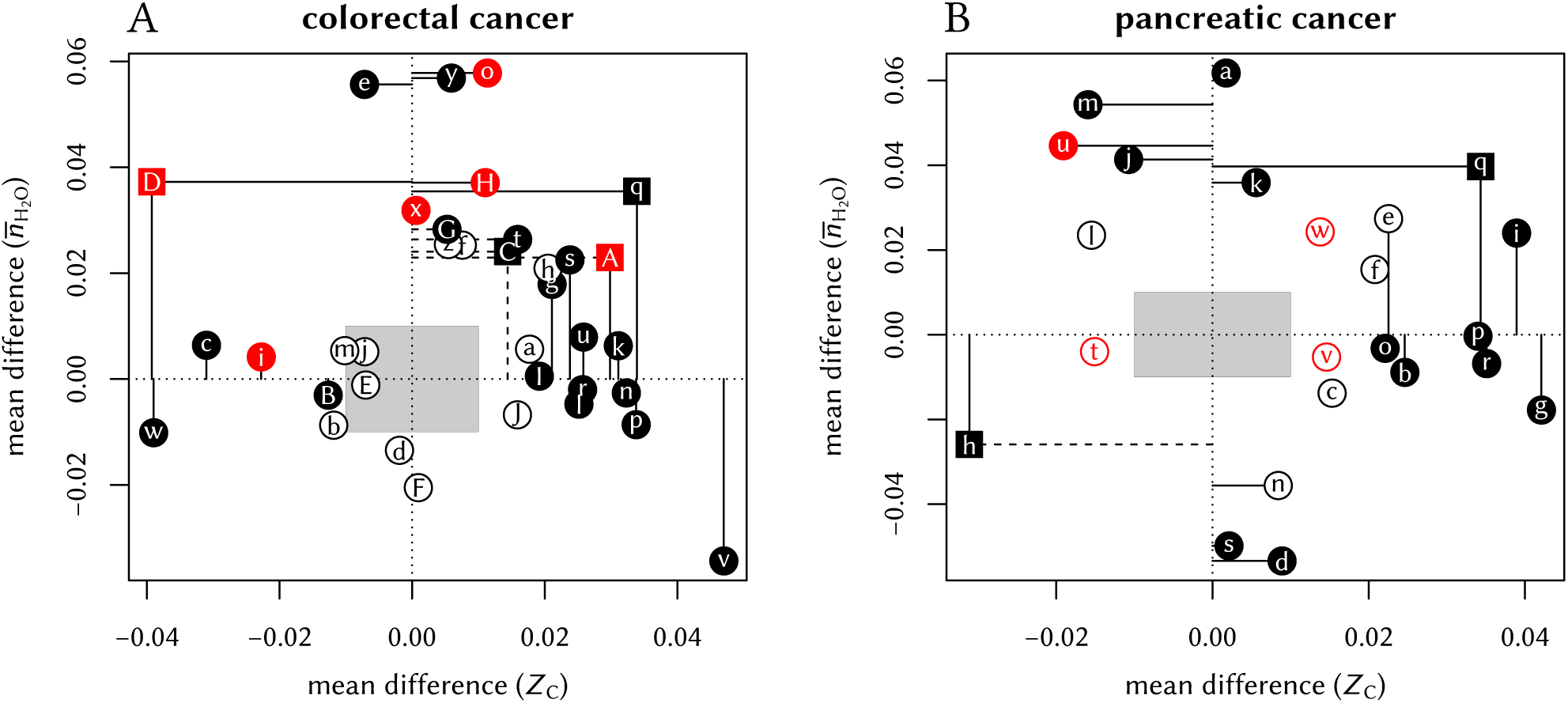
Compositional differences for protein expression in (A) colorectal cancer and (B) pancreatic cancer. Red colors highlight (A) adenoma / normal comparisons or (B) experiments in mice. Here and in Fig. 2, filled symbols and dashed lines indicate *p*<0.05; solid lines are drawn instead if the common language effect size is ≥60% or ≤40%.

The plot illustrates that proteins up-expressed in carcinoma relative to normal tissue most often have significantly higher *Z*_C_ [ⓖ ⓚ ⓛ ⓝ ⓟ ⓡ ⓢ ⓤ ⓥ ⓛ], *n*̅H_2_O [ⓔ ⓞ ⓣ ⓧ ⓨ Ⓓ Ⓖ Ⓗ], or both [ⓠ Ⓐ Ⓒ] (see also Dick, 2016). The red symbols in the plot highlight the datasets for adenoma / normal comparisons [ⓘ ⓞ ⑧ Ⓐ Ⓓ Ⓗ]. Most of these exhibit a significant positive Δ*n*̅H_2_O but not the large increase in *Z*_C_ found for many of the carcinoma / normal comparisons.

### Pancreatic cancer

Many proteomic studies have been performed to investigate the differences between normal pancreas (NP), early-stage pancreatic intraepithelial neoplasia, and pancreatic adenocarcinoma (PDAC). Proteomic studies also address the inflammatory condition of autoimmune pancreatitis, which is sometimes misidentified as carcinoma (Paulo et al., 2013), and chronic pancreatitis, which is associated with increased cancer risk (Chen et al., 2007). Searches for proteomic data were aided by the reviews of Pan et al. (2013) and Ansari et al. (2014). Table 2 lists selected datasets reporting at least ca. 30 down-expressed and 30 up-expressed proteins.

The compositional comparisons in Fig. 1B show that up-expressed proteins in pancreatic cancer often have significantly higher *Z*_C_ [ⓑ ⓔ ⓖ ⓘ ⓞ ⓟ ⓡ ⓡ].A dataset obtained for pancreatic cancer associated with diabetes mellitus (Wang et al., 2013a) [ⓠ] has both significantly higher *Z*_C_ and *n*̅H_2_O Only one dataset, from a study that targeted accessible proteins (Turtoi et al., 2011) [ⓗ], is characterized by a negative mean difference of *Z*_C_.Some other datasets that do not have significantly different *Z*_C_ exhibit either higher [ⓐ ⓙ ⓚ ⓜ ⓤ] or lower [ⓓ ⓗ ⓝ ⓢ] *n*̅H_2_O in cancer. Three of the four datasets with negative mean difference of *n*̅H_2_O are derived from particular study designs: chronic pancreatitis (Chen et al., 2007), accessible proteins (Turtoi et al., 2011), and low-grade tumors (Wang et al., 2013b). Therefore, the datasets with positive Δ*n*̅H_2_O may reflect a general characteristic of pancreatic cancer.

### Hypoxia and 3D culture

Hypoxia refers to oxygen concentrations that are lower than normal physiological levels. Hypoxia characterizes many pathological conditions, including altitude sickness, stroke, and cardiac ischemia. In tumors, irregular vascularization and abnormal perfusion contribute to the formation of hypoxic regions (Höckel and Vaupel, 2001). A related situation is the growth in the laboratory of 3D cell cultures (e.g. tumor spheroids), in contrast to two-dimensional growth on a surface. In 2D monolayers, all cells are exposed to the gas phase. In contrast, interior regions of 3D cultures are often diffusion-limited, leading to oxygen deprivation and necrosis (McMahon et al., 2012). There is some overlap, but also many differences, between gene expression in 3D culture and hypoxic conditions (DelNero et al., 2015).

Table 3 lists selected proteomic datasets with a minimum of ca. 20 down- and 20 up-expressed proteins in hypoxia or 3D growth. The differences in chemical composition of the differentially expressed proteins are plotted in Fig. 2A. In many datasets, hypoxia or 3D growth induces the formation of proteins with significant and/or large reduction in *Z*_C_ [ⓐ ⓑⓒ ⓖ ⓗ ⓙ ⓜ ⓞ ⓦ Ⓐ Ⓔ]. These datasets cluster around a narrow range of Δ*Z*_C_ (-0.032 to -0.021), except for dataset Ⓔ (3D growth of colon cancer cells) with much lower Δ*Z*_C_. As extracellular proteins tend to be more oxidized (Dick, 2014), the observation in some experiments that hypoxia decreases the abundance of proteins associated with the extracellular matrix (ECM) (Blankley et al., 2010) is compatible with the overall formation of more reduced (low-*Z*_C_) proteins. Conversely, reoxygenation leads to the formation of more oxidized proteins in supernatant (-S) and pellet (-P) fractions of isolated chromatin [ⓡ ⓤ].

**Figure 2.**
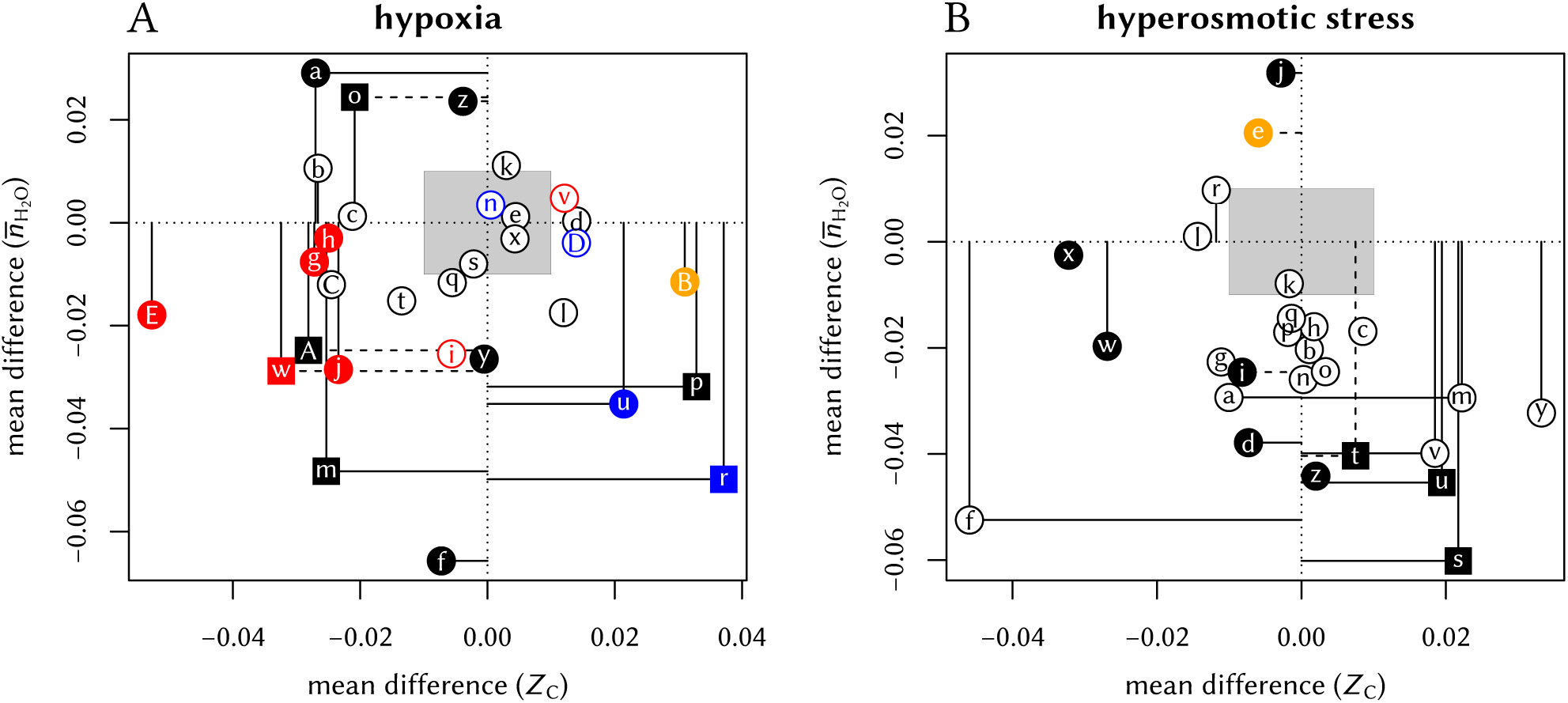
Compositional differences for protein expression in (**A**) hypoxia and (**B**) hyperosmotic stress. Red, blue, and orange symbols are used to highlight datasets for tumor spheres, reoxygenation or anti-hypoxic treatment, and adipose-derived stem cells, respectively.

While most studies controlled gas composition to generate hypoxia, two datasets [Ⓒ Ⓓ] are from a study that used cobalt chloride (CoCl_2_) to induce hypoxia in rat cardiomyocytes; treatment with salidroside (SAL) had anti-hypoxic effects (Xu et al., 2016). The CoCl_2_ and SAL treatments shift the proteome toward more reduced and more oxidized proteins, respectively, in agreement with the general trends for hypoxia and reoxygenation experiments.

Two datasets oppose the general trends, showing large and significantly higher *Z*_C_ under hypoxia. These datasets may be distinguished from the others by the particular conditions under which they were obtained. One of the nonconforming datasets is for the supernatant in a chromatin isolation procedure [ⓟ], and the other is for adipose-derived stem cells [Ⓑ],

### Hyperosmotic stress

By hyperosmotic stress is meant a condition that increases the extracellular hypertonicity, or osmolality. The addition of osmolytes (or “cosolvents”) lowers the water activity in the medium (Timasheff, 2002). Equilibration with hypertonic solutions drives water out of cells, causing cell shrinkage. The selected datasets listed in Table 4 include at least ca. 20 down-expressed and 20 up-expressed proteins in response to high concentrations of NaCl (5 studies), glucose (6 studies), succinate (1 study), KCl (1 study), or adaptation to seawater (1 study). The proteomic analyses used bacterial, yeast, or mammalian cells, or fish (eel) gills (Tse et al., 2013). One study varied temperature along with NaCl concentration (Kocharunchitt et al., 2012), and one study reported both transcriptomic and proteomic ratios (Kohler et al., 2015).

In the study of Giardina et al. (2014) [ⓞ ⓟ ⓠ], the reported expression ratios for transfer from low glucose to high glucose media are nearly all less than 1. Therefore, the “up-expressed” proteins in the comparisons here are taken to be those that have a higher expression ratio than the median in a given experiment. To achieve a sufficient sample size using data from Chen et al. (2015) [ⓡ], the comparisons here use a combined set of proteins, i.e. those identified to have the same direction of change in the two treatment conditions (380 and 480 mOsm NaCl) and a significant change in at least one of the conditions.

Fig. 2B shows that hyperosmotic stress strongly (CLES ≤40%) and/or significantly (*p*-value <0.05) induces the formation of proteins with relatively low water demand per residue in 11 datasets [ⓐ ⓑ ⓓ ⓕ ⓘ ⓜ ⓢ ⓣ ⓤ ⓥ ⓩ]. Five of these datasets, including 4 from bacteria [ⓢ ⓣ ⓤ ⓥ] and one from human cells [©], also show an increase in *Z*_C_. These trends are found in both the transcriptomic [ⓢ ⓣ] and proteomic [ⓤ ⓥ] data from the study of Kocharunchitt et al. (2012).

Four datasets obtained for mammalian cells have low Δ*Z*_c_ with no significant [ⓡ ⓦ ⓧ] or significantly negative mean difference of *n*̅H_2_O [ⓕ]. Six datasets [ⓗ ⓚ ⓝ ⓞ ⓟ ⓠ] from one study each in yeast and *E. coli*, and Japanese eels adapted to seawater, have very small mean differences in *Z*_C_ and a negative Δ*n*̅H_2_O that follows the trends of most of the other datasets, but with lower significance (*p*-value >0.05).

The comparisons show that hyperosmotic stress very often induces the formation of proteins with lower water demand per residue. In some, but not all, cases, this coincides with an increase in average oxidation state of carbon. Less often, and perhaps specific to mammalian cells, proteins are formed with a lower oxidation state of carbon. There is infrequent evidence, from two datasets with NaCl treatment [ⓔ ⓙ], for an increase in water demand per residue.

Notably, the two datasets for adipose-derived stem cells oppose the general trends for both hypoxic and hyperosmotic conditions (see Fig. 2A [Ⓑ] and Fig. 2B [ⓔ]). This intriguing result shows that these stem cells respond to external stresses with proteomic transformations that are chemically similar to those in cancer (Fig.1)

### Potential diagrams

Thermodynamic models can help to illuminate the microenvironmental context for proteomic differences. In particular, combining the chemical affinities of individual stoichiometric formation reactions of proteins yields an estimate of the thermodynamic potential for the overall process of proteomic transformation.

The chemical affinity quantifies the potential, or propensity, for a reaction to proceed. It is the infinitesimal change with respect to reaction progress of the negative of the Gibbs energy of the system. The chemical affinity is numerically equal to the “non-standard” or actual (Warn and Peters, 1996), “real” (Zhu and Anderson, 2002), or “overall” (Shock, 2009) negative Gibbs energy of reaction. These energies are not constant, but vary with the chemical potentials, or chemical activities, of species in the environment. Chemical activity (*a*) and potential (*μ*) are related through *μ* = *μ*° + *RT* ln *a,* while the standard chemical potentials (*μ*° = *G*°, i.e. standard Gibbs energies of species) are functions solely of temperature and pressure.

The equilibrium constant (*K*) for a reaction is given by Δ*G*° = −2.303*RT*log*K*, where Δ*G*° is the standard Gibbs energy of the reaction, 2.303 stands for the natural logarithm of 10, *R* is the gas constant, *T* is temperature in Kelvin, and log denotes the decadic logarithm. The equation used for affinity (***A***) is ***A*** = 2.303RT log(*K*/*Q*), where *Q* is the activity quotient of the reaction (e.g. Helgeson, 1979, Eq. 11.27; Warn and Peters, 1996, Eq. 7.14; Shock, 2009). Accordingly, the per-residue affinity of Reaction R1 can be written as

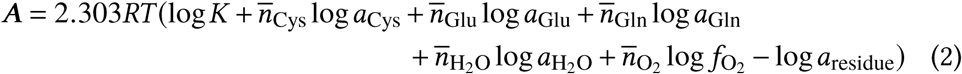

where we have substituted the abbreviations of the amino acids for their formulas. The overbar notation (*n*̅) signifies that the coefficients in Reaction R1 are each divided by the length (number of amino acids) of the protein sequence. Likewise, the elemental composition and standard Gibbs energy per residue are those of the protein (with formula C_*c*_H_*h*_N_*n*_O_*O*_S_*s*_) divided by the length of the protein.

The standard Gibbs energies of species at 37 °C and 1 bar were calculated with CHNOSZ (Dick, 2008) using equations and data taken from Wagman et al. (1982) and Kelley (1960) (O_2_(*_g_*), Johnson et al. (1992) and references therein (H_2_O), and using the Helgeson-Kirkham-Flowers equations of state (Helgeson et al., 1981) with data taken from Amend and Helgeson (1997) and Dick et al. (2006) (amino acids), and Dick et al. (2006) and LaRowe and Dick (2012) (amino acid group additivity for proteins).

Previously, activities of the amino acid basis species and protein residues (e.g. Eq.2) were set to 10^−4^ and 10^0^, respectively (Dick, 2016). As long as constant total activity of residues is assumed, the specific value does not greatly affect the calculations; here it is left unchanged at 10^0^. Revised activities of the amino acid basis species, corresponding to mean concentrations in human plasma (Tcherkas and Denisenko, 2001), are used here: 10^−3.6^ (cysteine), 10^-4.5^ (glutamic acid) and 10^−3.2^ (glutamine). Adopting these activities of basis species, instead of 10^−4^, lowers the calculated equipotential lines for proteomic transformations in hyperosmotic stress by about 0.5 to 1 log unit, closer to log *a*H_2_O = 0 (see below).

It follows from Eq. (2) that varying the fugacity of O_2_ and activity of H_2_O alters the chemical affinity for formation of proteins in a manner dependent on their chemical composition. For example, Figure 5A of Dick (2016) shows that increasing log*f*O_2_ is relatively more favorable for the formation of up-expressed than down-expressed proteins in a cancer dataset (Knol et al., 2014; ⓦ in Table 1). This tendency is consistent with the higher Zc of these up-expressed proteins (Fig. 1A).

How can we compare the affinities of groups, rather than individual proteins? One method is based on differences of the ranks of chemical affinities of proteins in two groups (Dick, 2016). Using this method, the affinities of all the proteins in a dataset are ranked; the ranks are then summed for proteins in the up- and down-expressed groups (*r*_up_ and *r*_down_). Before taking the difference, the ranks are multiplied by a weighting factor to account for the different numbers of proteins in the groups. This weighted rank difference (WRD) of affinity is a convenient summary of the differential potential for formation:

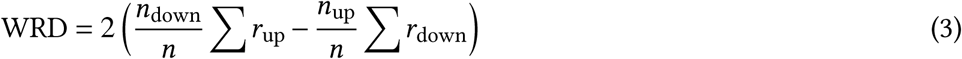

On a contour diagram of the WRD of affinity (“potential diagram”), the line of zero WRD represents a rank-wise equal potential (“equipotential”) for formation of proteins in the two groups.

To facilitate their interpretation, diagrams have been made for groups of proteomic datasets with similar compositional features. For pancreatic cancer, there are 11 datasets with Δ*Z*_C_ > 0.01 (i.e. to the right of the gray box in Fig. 1B) and for which the mean difference of *n*̅H_2_O is neither significant (low *p*-value) nor large (high CLES). Conversely, there are 8 datasets for pancreatic cancer with Δ*n*̅H_2_O > 0.01 and for which the mean difference of *Z*_C_ is neither large nor significant. Similarly, weighted rank-difference diagrams were constructed for 13 Δ*Z*_C_ > 0.01) and 10 (Δ*n*̅H_2_O > 0.01) datasets for CRC, 8 datasets for hypoxia Δ*Z*_C_ < −0.01), and 12 datasets for hyperosmotic stress (Δ*n*̅H_2_O < −0.01). The individual diagrams for each of these groups are presented in Figure S1.

In order to smooth out the variability between datasets and observe the central tendencies, the potential diagrams for each group in Figure S1 were combined by taking the arithmetic mean of the WRD at all grid points in log*f*O_2_ -log *a*H_2_O space. The resulting diagrams, in Fig. 3, have equipotential lines, shown in white, and zones of positive and negative WRD of affinity, i.e. greater relative potential for formation of up- and down-expressed groups of proteins, colored red and blue, respectively.

**Figure 3.**
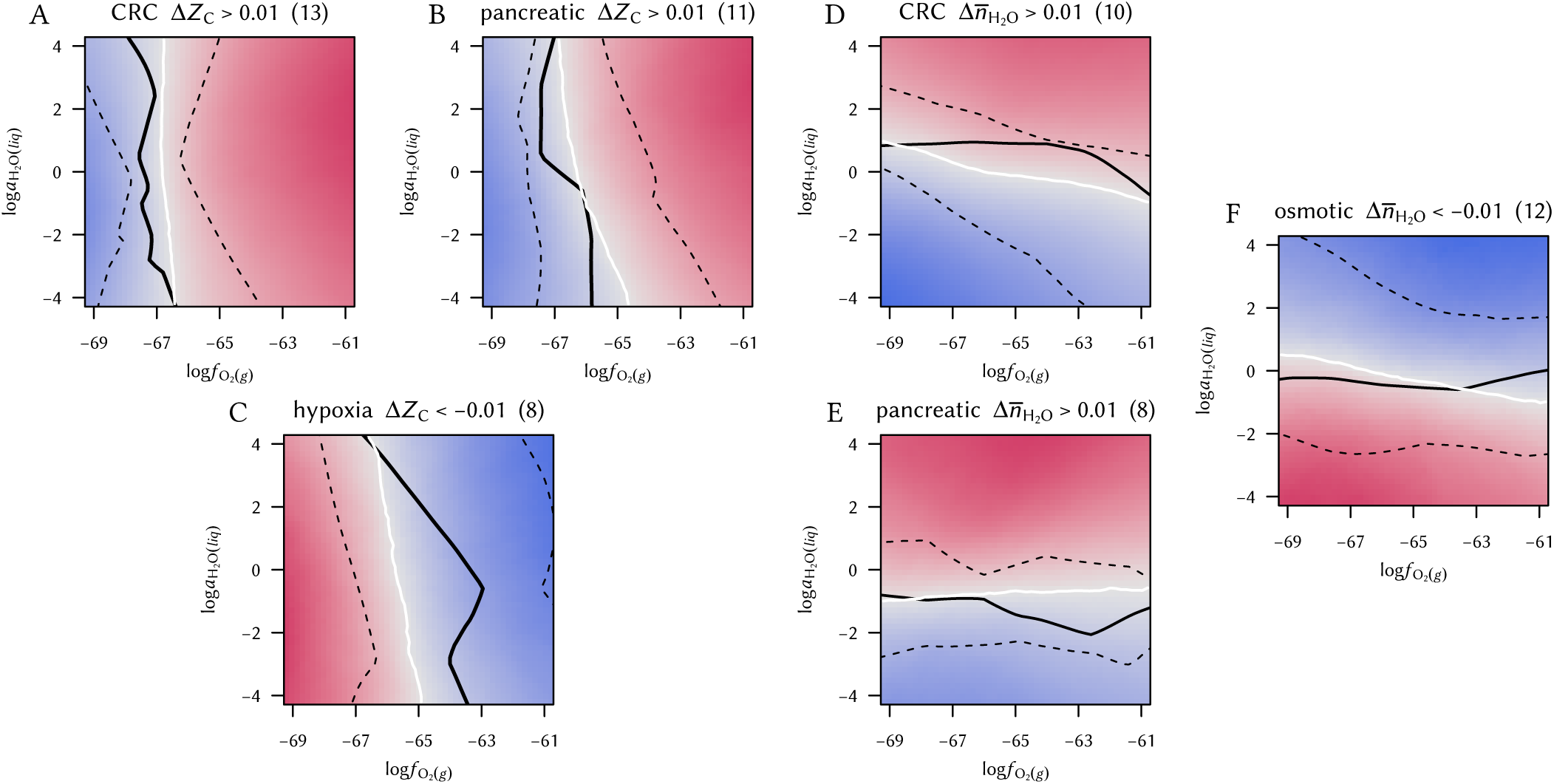
Merged potential diagrams for proteomic datasets. The groups consist of datasets with same-signed mean differences of *Z*_C_ or *n*̅H_2_O.See text for details.

The solid black lines in Fig. 3 show the median position along the *x*- or *y*-axis for the equipotential lines in each group (Figure S1), and the dashed black lines are positioned at the 1st and 3rd quartiles. The interquartile ranges for the cancer groups are smaller than those for hypoxia or hyperosmotic stress. This would be expected if the cancer datasets reflect a somewhat narrower range of conditions than the experimental datasets; the latter represent a wide a variety of organisms, cell types, and laboratory conditions (Tables 3–4).

## DISCUSSION

Calculations of the average oxidation state of carbon and water demand per residue, derived from elemental stoichiometry, shed new light on the microenvironmental context of differential protein expression in cancer and laboratory experiments. Hypoxia or hyperosmotic stress generally induces the formation of proteins with lower oxidation state of carbon or lower water demand per residue, respectively; here, the chemical makeup of the proteomes directly reflects the environmental stresses. In contrast, proteomes of CRC and pancreatic cancer are often characterized by greater water demand per residue or oxidation state of carbon. The formation of more highly oxidized proteins despite the hypoxic conditions of many tumors hints at a complex set of microenvironmental-cellular interactions in cancer.

Specific environmental ranges of oxygen fugacity and water activity affect the potential for proteomic transformation. The equipotential lines for cancer proteomes with high water demand and for proteomes in hyperosmotic stress lie close to log log *a*H_2_O = 0 (Fig. 3D-F). Although there is considerable variability among the individual datasets (Figure S1; see also Dick, 2016), the merged diagrams demonstrate a physiologically realistic range for the activity of water. Thus, the ensemble of lower-level processes regulating proteomic transformations likely operate in conjunction with the chemical constraints of an aqueous environment. Water activity in cells is close to one, but restricted diffusion of H_2_O in “osmotically inactive” regions of cells (Model, 2014) could result in locally lower water activities, and may account for some of the variability in the datasets.

The finding of a frequently positive water demand for the transformation between normal and cancer proteomes offers a new perspective on the biochemistry of hydration in cancer. The thermodynamic calculations predict that, in contrast to hyperosmotic stress, proteomes of cancer tissues are stabilized by water activity ≳ 1. A generally higher water activity would be consistent with an increased hydration of tissue, which is apparent in spectroscopic analysis of breast cancer (e.g. Abramczyk et al., 2014). Speculatively, the relatively high water content needed for embryonic development (Moulton, 1923) could be recreated in cancer cells if they revert to an embryonic mode of growth (McIntyre, 2006).

The equipotentials for transformation of proteomes in cancer cluster near an oxygen fugacity of 10^−66^. The oxygen fugacity should be interpreted not as actual oxygen concentration, rather as a scale of the internal oxidation state of the system. The oxygen fugacity and water activity can be converted to the Eh redox scale, giving values that are comparable to biochemical measurements (Dick, 2016).

Although cancer proteomes are derived from cells that are often exposed to hypoxic tumor environments, the differential expression is most often in favor of oxidized proteins (Fig. 1A and B). What are some explanations for this finding? Perhaps the relatively high log *f*o_2_ threshold for chemical transformation of hypoxia-responsive proteins could support a buffering action that potentiates the formation of relatively oxidized proteins in cancer (compare the median and upper quartile in Fig. 3C with those in Fig. 3A and B). This speculative hypothesis requires a division of the cellular proteome into localized, chemically interacting subsystems. Alternatively, the development of a high oxidation potential in cancer cells may be associated with a higher concentration of mitochondrially produced reaction oxygen species (ROS). Again, this suggests that a physical separation of cellular macromolecules is responsible for the observed proteomic differences.

Neither of these possibilities addresses the magnitude of the chemical differences in the proteomes, and the question remains: where do the electrons go?

One hypothesis comes from the different oxidation states of biomolecules. Fatty acids are highly reduced compared to amino acids, nucleotides, and saccharides (e.g. Amend et al., 2013). In parallel with the formation of more reduced proteins, hypoxia induces the accumulation of lipids in cell culture (Gordon et al., 1977). Cancer cells are also known for increased lipid synthesis. Lipid droplets, which are derived from the endoplasmic reticulum (ER), form in great quantities in cancer cells (Koizume and Miyagi, 2016). Assuming that lipids are synthesized from relatively oxidized metabolic precursors, their formation requires a source of electrons. These considerations lead to the hypothesis that lipid synthesis is coupled, through the transfer of electrons, to the oxidation of the proteome.

Here, calculations that combine proteomic and cellular data are used to quantify a hypothetical redox balance among pools of lipids and proteins. The major assumptions in this calculation are that the overall cellular oxidation state of carbon is the same in cancer and hypoxia, and that changes in this cellular oxidation state are brought about by altering only the numbers of lipid and protein molecules. The overall chemical composition of the lipids is assumed to be constant, but the proteins are assigned different values of *Z*_C_. These simplifying assumptions are meant to pose quantifiable “what if” questions, to serve as points of reference about the range of molecular composition of cells (Milo and Phillips, 2015).

The worked-out calculation is shown in Fig. 4. The lipid:protein ratio in hypoxia is taken from Gordon et al. (1977), and ballpark values for the differences in Zc of proteins in hypoxia and cancer are from the present study. Notably, the lipid:protein weight ratio in hypoxia (0.19) is higher than in normal cells (i.e. 0.15 using data from Gordon et al., 1977 or 0.16 using data compiled by Milo and Phillips, 2015 for *E. coli*).

**Figure 4.**
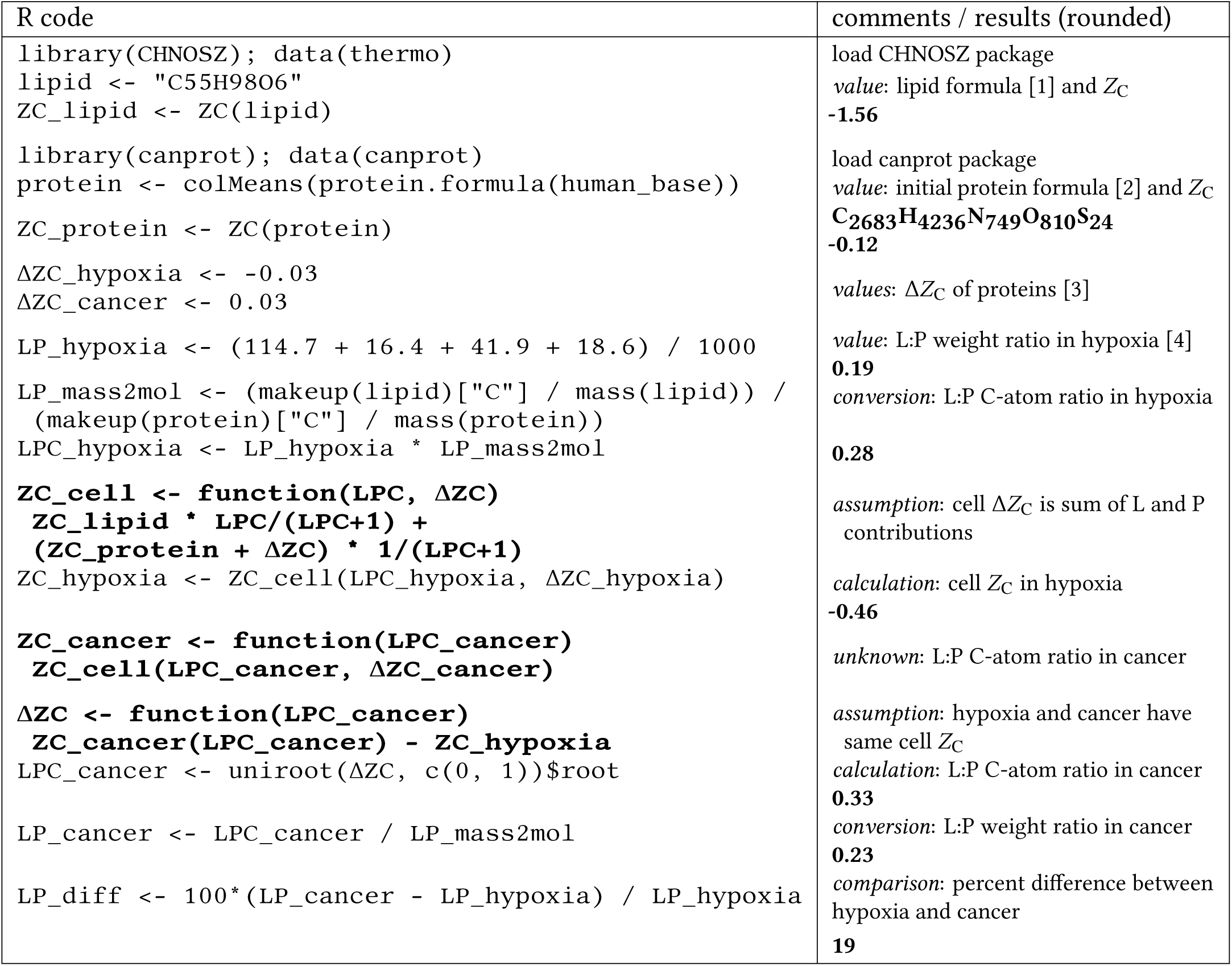
A computer-aided “back of the envelope” calculation to estimate the lipid to protein ratio (L:P) in cancer cells and the percent difference from normal cells in hypoxic conditions.Bold text indicates function definitions (left column) or numerical results (right column). Numerical values are taken from [1] Wikipedia: triglyceride, [2] UniProt: human proteome (human_base.Rdata in the **canprot** package), [3] this study, and [4] Gordon et al. (1977) (mouse cells grown in hypoxic conditions).

The calculation indicates that an increase of the lipid:protein weight ratio in cancer cells by ca. 20% over that in hypoxic normal cells could provide an electron sink that is sufficient to take up the electrons released by oxidation of the proteome in cancer Δ*Z*_C_ ≈ 0.03) compared to hypoxic normal cells Δ*Z*_C_ ≈ −0.03 relative to to non-hypoxic normal cells). This result shows a predicted higher lipid:protein ratio in cancer cells or tissues than normal cells under hypoxic conditions.

As found by Raman spectroscopy, levels of both lipids and proteins are elevated in colorectal cancer (Stone et al., 2004), while CRC stem cells can have a higher lipid:protein ratio than cancer and normal epithelial cells (Tirinato et al., 2015). In contrast to CRC, lipids are decreased in breast cancer compared to normal breast tissue (Frank et al., 1995; Stone et al., 2004). Given a lower lipid content, and therefore smaller electron sink, one might expect that proteomes in breast cancer are oxidized to a lesser extent than those in CRC and pancreatic cancer. Preliminary analysis of proteomic data for breast cancer (not shown) support this prediction. Other factors that affect the systematic redox balance, such as a more reduced gut microbiome in CRC (Dick, 2016) and metabolic coupling between epithelial and stromal cells, may be important for an accurate account of the compositional biology.

These compositional and thermodynamic analyses support the notion that physical constraints imposed by microenvironmental conditions have a primary role in shaping the relative abundances of proteins. This approach to the data differs from conventional interpretations of proteomic data based on functions of proteins. However, the scope of explanations dealing with functions and molecular interactions offers limited insight on the high-level organization of proteomes in a cellular and microenvironmental context. Although a variety of bioinformatics tools are available for functional interpretations (Laukens et al., 2015), none so far addresses the overall chemical requirements of proteomic transformations. The compositional and thermodynamic descriptions presented here encourage a fresh look at the question, “What is cancer made of?”

## CONCLUSION

Although many hypoxia experiments induce the formation of proteins with lower oxidation state of carbon (*Z*_C_), proteomes of colorectal and pancreatic cancer are often relatively oxidized compared to normal tissue. Hyperosmotic stress in the laboratory generally leads to the formation of proteins with relatively low water demand per residue (*n*̅H_2_O), but cancer proteomes often show the opposite trend.

The global proteomic differences can be described and quantified within a componential thermodynamic framework. This high-level perspective uses concepts and language that differ from those applicable to molecular interactions (Ellis, 2015). A positive thermodynamic potential for any proteomic transformation is achieved in a specific, physiologically relevant range of oxidation and hydration potential. Speculatively, further application of these methods could be used to predict the ability of chemotherapy or other treatments to reduce or reverse the potential for formation of the proteins required by cancer cells. Findings of low oxidation and/or hydration potential in a thermodynamic analysis of differential protein expression are likely to be characteristic of beneficial treatments.

The distribution of biomolecules other than proteins should be considered to account for changes in cellular redox balance. An electron sink associated with a ca. 20% greater lipid to protein ratio in cancer compared to normal hypoxic cells would be sufficient to balance the electrons released by the formation of more oxidized proteins in CRC and pancreatic cancer. It thus appears possible that a redox disproportionation develops in some cancers, leading to pools of both more reduced and more oxidized molecules compared to normal conditions.

## ACKNOWLEDGEMENTS

I thank Apar Prasad for commenting on the paper and Dr. Ming-Chih Lai for providing data generated in the study of Lai et al. (2016) and giving permission to include it here.

## SUPPLEMENTAL INFORMATION

### Dataset S1

R source package including amino acid composition and protein expression data (canprot_0.0.3.tar.gz).

### Dataset S2

Project code file.

### Table S1

Compositional summaries: mean values of *Z*_C_ and *n*̅H_2_O and corresponding mean differences, *p*-values, and common-language effect sizes (CLES).

### Figure S1

Potential diagrams for individual datasets. These diagrams were merged to make the diagrams in Fig. 3.

